# Knocking out alpha-synuclein in melanoma cells downregulates L1CAM and decreases motility

**DOI:** 10.1101/2023.01.17.524111

**Authors:** Nithya Gajendran, Santhanasabapathy Rajasekaran, Stephan N. Witt

**Author notes:** equal contributions (Nithya Gajendran, Santhanasabapathy Rajasekaran).

## Abstract

The Parkinson’s disease (PD) associated protein, alpha-synuclein (α-syn/*SNCA*), is highly expressed in aggressive melanomas, which raises the possibility that α-syn has a pro-survival function in melanoma. Herein, we asked whether α-syn modulates the expression of the pro-oncogenic adhesion molecules L1CAM and N-cadherin. We used two human melanoma cell lines (SK-MEL-28, SK-MEL-29), *SNCA*-knockout (KO) clones, and two human SH-SY5Y neuroblastoma cell lines. In the melanoma lines, loss of α-syn expression resulted in significant decreases in the expression of L1CAM and N-cadherin and concomitant significant decreases in motility. On average, there was a 75% reduction in motility in the four *SNCA*-KOs tested compared to control cells. Strikingly, comparing neuroblastoma SH-SY5Y cells that have no detectable α-syn to SH-SY5Y cells that stably express α-syn (SH/+αS), we found that expressing α-syn increased L1CAM and single-cell motility by 54% and 597%, respectively. The reduction in L1CAM level in *SNCA*-KO clones was not due to a transcriptional effect, rather we found that L1CAM is more efficiently degraded in the lysosome in *SNCA*-KO clones than in control cells. We propose that α-syn is pro-survival to melanoma (and possibly neuroblastoma) because it promotes the intracellular trafficking of L1CAM.

## Introduction

Melanoma is an aggressive skin cancer that arises from pigment-producing cells called melanocytes. Epidemiological studies have reported the co-occurrence of melanoma and Parkinson’s disease (PD)^1,2^. Such a co-occurrence is unusual in that these two diseases are so different in that melanoma is characterized by uncontrolled cell proliferation whereas PD by neuronal cell death. Melanoma patients have a 1.5- to 1.85-fold higher risk of developing PD compared to age- and sex-matched controls^3,4^, and reciprocally, PD patients have a 1.4- to 20-fold higher risk of developing invasive melanoma than a control group^5-7^. The mechanism of this co-occurrence is unknown and could involve multiple genes^2^, one of which is *SNCA*, and even environmental factors. Here we focus on the role of *SNCA* in melanoma.

*SNCA* codes for the protein alpha-synuclein (α-syn)^8-10^, which is expressed in dopaminergic neurons^11-13^ and a variety of other tissues^14-16^, including melanomas^17^, where it is highly expressed. α-Syn is a small intrinsically unfolded protein that in neurons localizes to nerve terminals where synaptic vesicles are docked; it accelerates the kinetics of individual exocytotic events^18^. Whether α-syn promotes exocytosis in other cell types is not known. Many other activities have been ascribed to this unique protein because it, in its multitude of conformations, binds to so many biomolecules, i.e., negatively charged phospholipids^19,20^, proteins^21-23^, and DNA^24^. Thus, while α-syn functions in the endolysosomal system to promote endocytosis and exocytosis^18,25^, and it may have other functions. The intrinsically unfolded nature of α-syn makes it highly prone to aggregate into oligomeric and prion-like, amyloidogenic conformations^26^, and some of these aggregates trigger neurodegeneration in PD^27^. Paradoxically, prion-like aggregates of α-syn have been suggested to promote autophagy in melanoma cells^28^, which is pro-survival. Whether α-syn has other pro-survival functions in melanoma is the subject of this study.

Two studies assessed the effect of knocking out *SNCA* on cellular homeostasis. First, knocking down *SNCA* in mouse retinal epithelial cells in vitro significantly decreased the level of the transferrin receptor (TfR1) and its mRNA transcript, and overexpressing α-syn increased the levels of the TfR1 protein and its mRNA transcript relative to control cells^29,30^. The loss of α-syn expression in retinal epithelial cells caused TfR1 molecules to accumulate in Golgi vesicles, suggesting that α-syn is required for the pathway that transits TfR1 from the *trans*-Golgi to the plasma membrane^29^. Second, *SNCA* was knocked out in the human cutaneous melanoma cell line, SK-Mel-28, and the *SNCA*-KO clones were assessed in vitro and in a mouse xenograft model^30^. The loss of α-syn expression in these melanoma cells decreased the levels of TfR1 and the iron exporter ferroportin and significantly suppressed the growth of the *SNCA-*KO tumors engrafted in nude mice. The reduction in the level of TfR1 in the *SNCA*-KO clones was a consequence of its enhanced lysosomal degradation. The results from these two studies are consistent with α-syn promoting the vesicular trafficking of TfR1.

In this study, we sought to determine whether the expression level of α-syn affects the expression level of adhesions proteins. Specifically, given that TfR1 and L1CAM, which is an adhesion protein expressed in neurons and melanomas, co-localize upon endocytosis in 3T3 cells^31^, we asked whether α-syn modulates the expression of L1CAM, like it does TfR1. To this end, we measured the levels of L1CAM in melanoma and neuroblastoma cells with or without the expression of α-syn. In addition, we also assessed the levels of E- and N-cadherin, and vimentin, which are three proteins involved in the epithelial-to-mesenchymal transition (EMT)^32,33^. The EMT is a highly regulated transition where epithelial cells shed their epithelial markers and morphology and convert to a mesenchymal phenotype. However, because melanocytes (and melanomas) are not epithelial cells, it is a misnomer to say that melanocytes undergo an EMT. We show that knocking out α-syn in melanoma cell lines and low expression of α-syn in a neuroblastoma cell line cause significant decreases in L1CAM compared to control cells that express α-syn. Our interpretation of these findings is that α-syn is a pro-survival factor in melanoma because it acts post-translationally to maintain a high levels of L1CAM and in turn high levels of motility.

## Results

The cell lines used in this study are given in Table 1. Each melanoma cell line harbors the BRAF V600E mutation^34^, which is the most common mutation in cutaneous melanoma (Table 1). This mutation causes constitutive activation of the RAS-RAF-MEK-ERK signaling pathway^35^, which leads to proliferation. SH-SY5Y cells, which are widely used to study PD^36^, are derived from a neuroblastoma.

**Table 1.**

| Cell line | Comments | α-syn (Y,N) | Mutations | ref |
| --- | --- | --- | --- | --- |
| SK-MEL-28 | This cell line was isolated from skin tissue from a 51-year-old, male with malignant melanoma. | Y | BRAF V600E | 17 |
| SNCA-KO clones KO8, KO9 | SNCA knocked out in SK-MEL-28 cells using CRISPR/Cas9. | N | BRAF V600E | 30 |
| SNCA-KI clones KI8, KI9 | SNCA cDNA was re-introduced into the KO clones using lenti virus. | Y | BRAF V600E | 30 |
| SK-MEL-29 | This cell line was isolated from a recurrent melanoma at the apex of the left axilla of a 19-year-old male. | Y | BRAF V600E |  |
| SNCA-KO clones KO1, KO2 | SNCA knocked out in SK-MEL-29 cells using CRISPR/Cas9. | N | BRAF V600E | This study |
| SH-SY5Y | Cell line derived from a neuroblastoma from a 4-year-old child. | N | unknown | 55 |
| SH/aS | SH-SY5Y stably expressing α-syn. | Y | unknown | 55 |

### SK-MEL-28 *SNCA*-KO cells have decreased levels of EMT-like markers and decreased invasion, migration, and motility

We first examined SK-MEL-28 control cells and their derivatives (Table 1). The levels of E-cadherin, L1CAM, N-cadherin, vimentin, α-syn, and α-tubulin were probed in cell extracts using Western blotting (Fig. 1A-D; Fig. S1, uncropped blots). The normalized band intensities from a densitometric analysis of the blots are shown in Figure 1E. Compared to the SK-MEL-28 control cells, the loss of α-syn expression significantly decreased the expression of L1CAM, N-cadherin, and vimentin in the two *SNCA*-KO clones by an average of 26% (*P*=0.002), 35% (*P*=0.005), and 52% (*P*=0.008), respectively. In contrast, the level of E-cadherin was unaffected (Fig. 1E). Clones KI8 and KI9 exhibited levels of these three proteins like the control cells.

**Figure 1.**
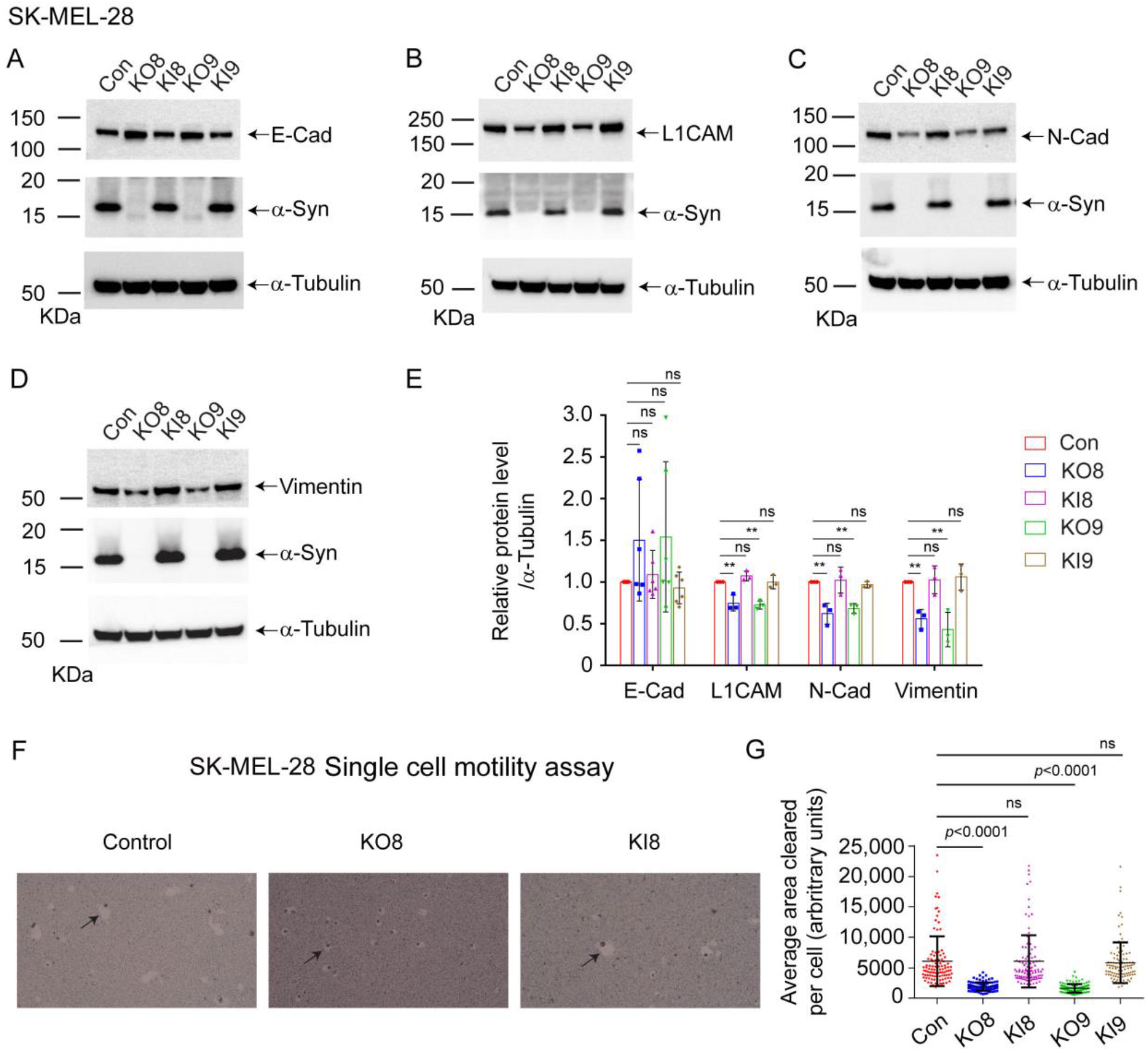
Loss of α-syn decreases EMT-like markers and motility in SK-MEL-28 cells. **(A-D**) Representative western blots of E-Cad, L1CAM, N-Cad, vimentin, α-syn, and α-tubulin in lysates of the control, KO, and KI cells cultured *in vitro*. (**E**) Quantitative analysis of relative protein levels. E-Cad, L1CAM, N-Cad, and vimentin band intensities are normalized to α-tubulin. All the experiments were repeated with at least three biological replicates (n=3). (**F**) Representative light microscope images of phagokinetic tracks created by control, KO8, and KI8 cells on colloidal gold-coated wells. Black arrows mark individual phagokinetic tracks in respective cell lines. Images were acquired by using a 10x objective, and the cleared phagokinetic area per cell was measured using ImageJ software. (**G**) Bar graphs show quantification of the cleared areas from three independent experiments. At least 33 tracks per experimental condition were randomly chosen for quantification. 100 tracks for n=3 per experimental group were used for statistical analysis. (**E, G**) Values are mean ± s.d. **, *p* = 0.0015 - 0.0027; *, *p* = 0.0073 - 0.0134 determined using a one-way ANOVA, Dunnett post hoc test.

Given that the *SNCA*-KO clones express lower levels of L1CAM, N-cadherin, and vimentin, which are linked to invasion and migration, we asked whether the KO clones are less motile than control cells. To this end, we used a colloidal gold single-cell motility assay^37^ to monitor the movement of single cells. As a cell moves across the surface of a slide covered with colloidal gold, the cell leaves a track where there is less gold on the surface. Representative images of the movement of control, KO8, and KI8 cells are shown in Figure 1F. Representative images of the movement of control, KO9, KI9 are shown in Figure S2A. The control cells created much larger tracks than the KO8 clones, and the KI8 clone pattern of tracks was like that of the control cells. ImageJ was used to measure the area of the tracks, and the results are plotted in Figure 1G. Loss of α-syn expression significantly decreased single-cell motility of KO8 and KO9 by 69% (*P*<0.0001) and 73% (*P*<0.0001), respectively, and re-expression of α-syn restored motility to the level of the control cells.

The invasion and migration potential of SK-MEL-28 control, *SNCA*-KO, and *SNCA*-KI cells were determined by transwell assays using Boyden chambers. The invasion assay measures the ability of cells in serum-free media in the top chamber to invade the matrigel and move into the lower chamber with complete media containing FBS. The migration assay uses the same setup but without matrigel. After 24 h, cells in the lower chamber were fixed, stained with crystal violet, imaged by light microscopy, and counted. Representative images from the migration assay that compared control, KO8, and KI8 are shown in Figure 2A; images for control, KO9, and KI9 are shown in Figure S2B. The number of migrated cells per field showed a significant 77% (*P*<0.0001) decrease for KO8 cells compared to the control cells, and KI8 exhibited the same level of migrated cells per field as the control cells (Fig. 2A, left-hand plot). Similar results were found for the KO9 and KI9 clones (Fig. 2A, right-hand plot). Representative images from the invasion assay that compared control, KO8, and KI8 are shown in Figure 2B; images for control, KO9, and KI9 are shown in Figure S2C. The number of invaded cells per field showed a significant 77% (*P*<0.0001) decrease for KO8 cells compared to control cells, and KI8 exhibited the same level of migrated cells as the control cells (Fig. 2B, left-hand plot). Similar results were obtained for the KO9 and KI9 clones (Fig. 2B, right-hand plot).

**Figure 2.**
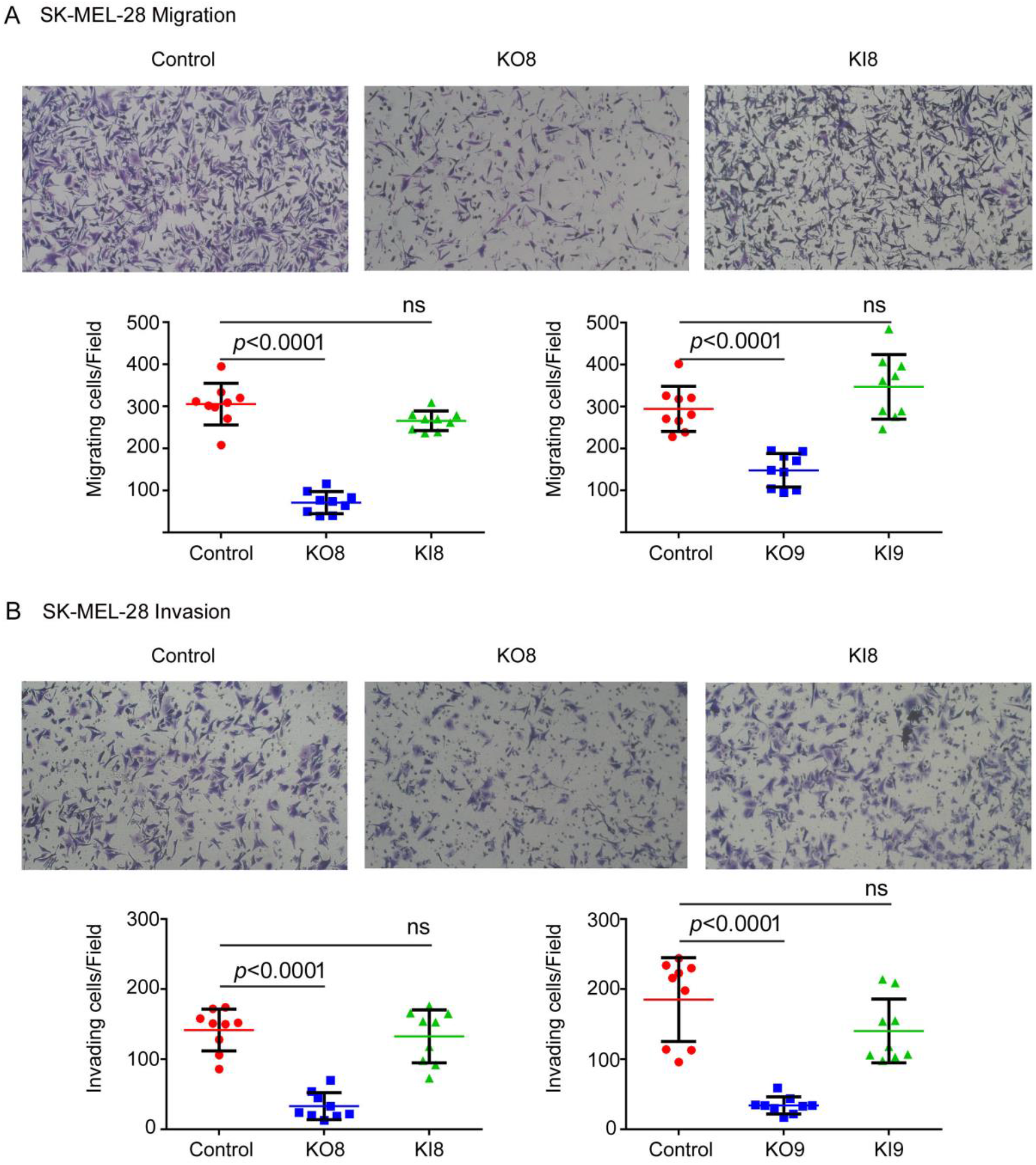
*SNCA-KO* reduces migration and invasion in SK-MEL-28 cells as assessed by transwell chamber assay. In the migration assay, 1 × 10^5^ cells in serum-free media were seeded in the upper chamber of the transwell (8 μm pore) apparatus and allowed to migrate through the membrane into the bottom chamber. In the invasion assay, 50 μl matrigel (0.2 mg/ml) was added to form a thin gel layer before the assay, and then 4 × 10^5^ cells in serum-free media were seeded in the upper chamber and allowed to invade through the membrane into the bottom chamber. In both experiments, DMEM complete media with 50% FBS in the lower chamber served as chemoattractant. Cells that passed through the membrane were fixed on the membrane with paraformaldehyde and stained with crystal violet. **(A**) Representative images of migrated control, KO8, and KI8 cells (after 24 hours) per 10X field. A total of three microscopic fields were randomly selected from each inner membrane and cells were counted and represented in graph (n=3 independent experiments). (**B**) Representative images of invaded control, KO8, and KI8 cells (after 24 hours) per 10X field. A total of three microscopic fields were randomly selected from each inner membrane, and cells were counted and represented in graphs (n=3 independent experiments). Values are mean ± s.d. *P*-values determined by a one-way ANOVA, Dunnett post hoc test.

Next, we asked why the level of the L1CAM protein is significantly reduced in *SNCA*-KO cells relative to control cells. We focused on L1CAM for the following reasons: (i) L1CAM and α-syn has been implicated in synaptic plasticity^38^. (ii) L1CAM is recognized as a tumor antigen involved in motility^39-41^. (iii) L1CAM is endocytosed with TfR1 in 3T3 fibroblast cells^31^. (iv) TfR1 molecules are more efficiently degraded in the lysosome in *SNCA*-KO cells compared to control cells^30^. L1CAM molecules might be more rapidly degraded in the absence of α-syn expression, or L1CAM mRNA might be downregulated in the absence of α-syn expression. Both possibilities were checked.

We tested for degradation of L1CAM molecules in the lysosome using the inhibitor bafilomycin A1 (baf). If L1CAM is degraded in the lysosome, baf will block its degradation; thus, the level of L1CAM in baf-treated *SNCA*-KO cells should be the same as in control cells. To this end, cells were treated for 5 h with baf (or dimethyl-sulfoxide, DMSO, as a control). The extracts were probed for L1CAM, LC3-I, II, α-syn, and α-tubulin by Western blotting (Fig. 3A; Fig. S3, uncropped blots). LC3-II, which is a lipidated form of LC3, builds up in autophagosomes when autophagy is inhibited. In the DMSO group, the expression of L1CAM in each of the *SNCA*-KO clones decreased on average 50% (*P* = 0.022-0.024) relative to control cells (Fig. 3A, lanes 2 and 4 vs 1; Fig. 3B); whereas, in the baf group, L1CAM showed no significant decrease in the KO clones relative to control cells (Fig. 3A, lanes 8 and 10 vs 7; Fig. 3B).

**Figure 3.**
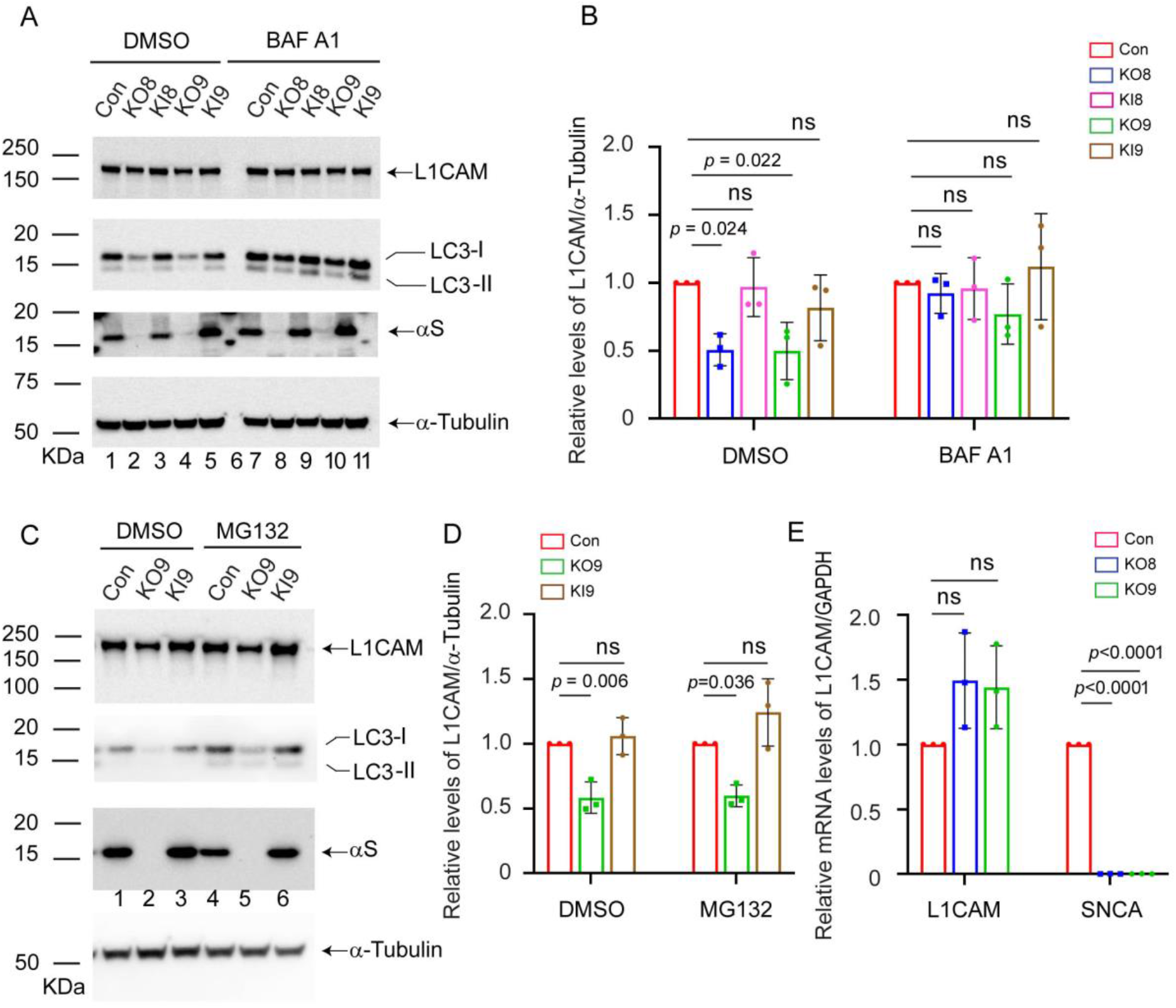
Loss of α-syn promotes the lysosomal degradation of L1CAM. (**A**) Representative Western blots showing the effect of bafilomycin A1 (baf) on the levels of L1CAM, LC3-II, and α-syn in lysates of the control, KO and KI cells cultured *in vitro*. Indicated cells were treated with 50 nM baf for 5 h and the lysates were probed for the indicated proteins. (**B**) Quantitative data showing the relative protein levels of L1CAM normalized to α-tubulin. (**C**) Representative Western blots showing the effect of MG132 on the levels of L1CAM, LC3-II, and α-syn in lysates of the control, KO9, and KI9 cells cultured *in vitro*. The cells were treated with 10 μM MG132 for 6 h and the lysates were probed for the indicated proteins. (**D**) Quantitative data showing the relative protein levels of L1CAM. The band intensity of L1CAM was normalized to α-tubulin. (**E**) Quantitative RT-PCR analysis of L1CAM. Relative mRNA levels in fold-change of L1CAM and *SNCA* normalized to housekeeping gene GAPDH. Values are mean ± s.d. *P*-values determined by a one-way ANOVA, Dunnett post hoc (n = 3).

We tested for degradation of L1CAM molecules by the proteasome using the inhibitor MG132. If L1CAM molecules are degraded in part by the proteasome, then the level of L1CAM in MG132-treated *SNCA*-KO cells should be the same as in control cells. To this end, cells were treated for 6 h with MG132 (or dimethyl-sulfoxide, DMSO, as a control), and the extracts were probed for L1CAM, α-syn, LC3-I, -II, and α-tubulin by Western blotting (Fig. 3C; Fig. S3, uncropped blots). However, MG132 did not increase the level of L1CAM in the KO9 cells, that is, in both groups L1CAM was 40% lower in the KO9 clone than in control cells (Fig. 3C, compare lanes 1 and 2 and 4 and 5; Fig. 3D). The results show that the decreased level of L1CAM in the *SNCA*-KO cells is due to the protein being degraded in the lysosome.

Quantitative PCR (qPCR) was conducted to determine whether the level of L1CAM mRNA was decreased in the *SNCA*-KO9 clone. No evidence was found for a decrease in L1CAM mRNA in KO9 cells compared to the control cells. Instead, a modest increase of this transcript in the KO9 clone was detected (Fig. 3E). Collectively, the data in Figures 1-3 show that loss of α-syn expression in SK-Mel-28 cells significantly decreases the levels of several proteins involved in invasion and migration; the decrease in the expression of L1CAM is due to lysosomal degradation; and the KO clones displayed significant decrements in migration, invasion, and motility relative to control cells.

### SK-MEL-29 *SNCA*-KO cells have decreased levels of EMT-like markers and decreased motility

We knocked out *SNCA* in the human cutaneous cell line SK-MEL-29 and used the parental cells and two *SNCA*-KO clones (Table 1) in the following experiments. Western blotting was used to probe the expression of L1CAM, N-cadherin, vimentin, α-syn, and α-tubulin in cell extracts (Fig. 4A, B; Fig. S4, uncropped blots). Loss of α-syn expression caused a 20% decrease in the expression of L1CAM in both *SNCA*-KO clones compared to control cells (Fig. 4 A, C), a 20% decrease in N-cadherin (Fig. 4A, D), but no change in vimentin (Fig. 4B, E). Although these decreases in protein expression were modest, nevertheless, three of the four changes were statistically significant. We also conducted the colloidal gold single-cell motility assay on SK-MEL-29 parental cells and two *SNCA*-KO clones. The control cells had much greater motility than either of the two KO clones, and images of the tracks for control and KO1 are shown in Figure 4 F.

**Figure 4.**
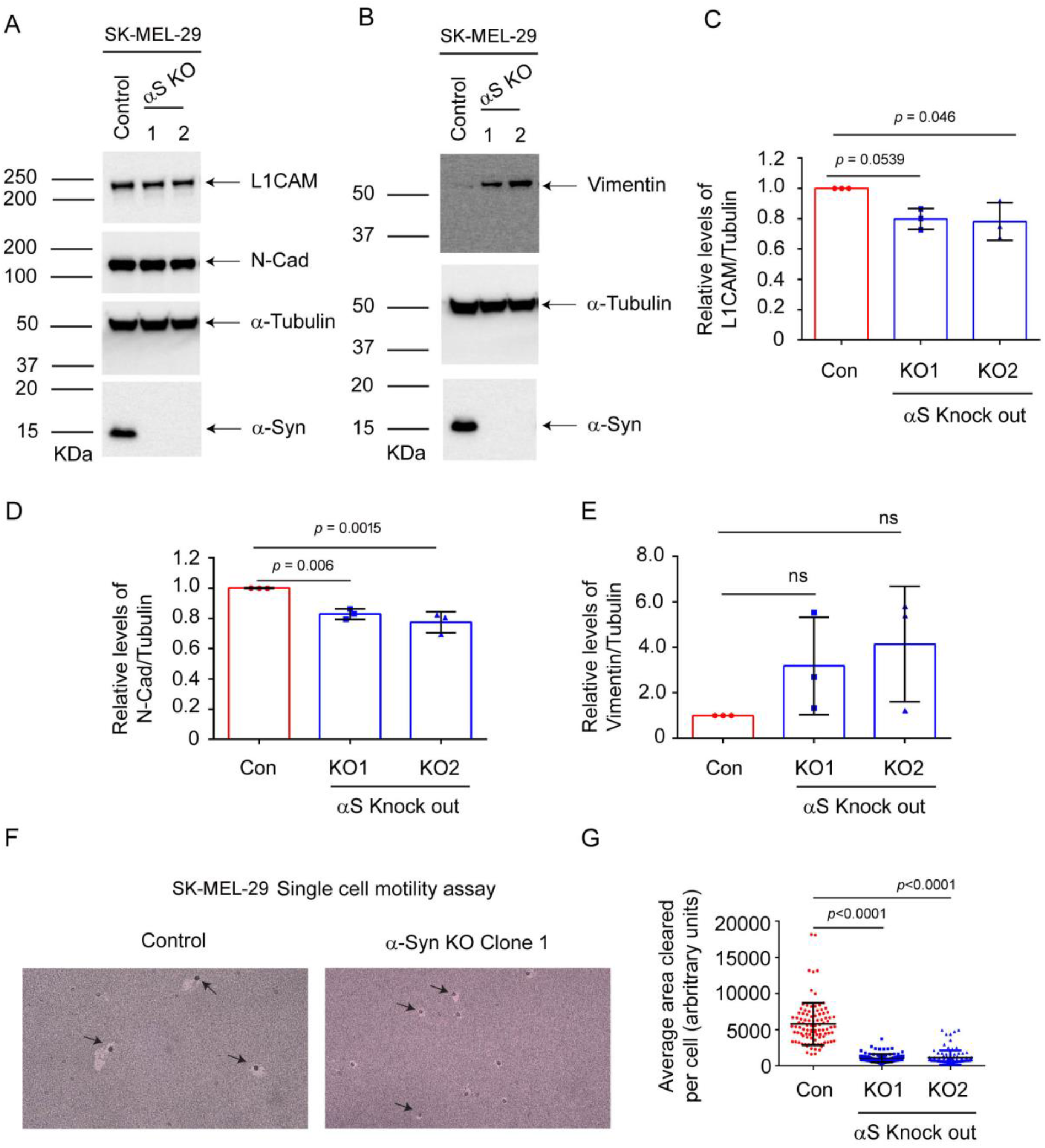
Loss of α-syn decreases L1CAM and N-cadherin and motility in SK-MEL-29 cells. (**A**) Representative Western blots of L1CAM, N-Cad, α-syn, and α-tubulin in lysates of the control and α-syn KO cells cultured *in vitro*. (**B**) Representative Western blots of vimentin, α-syn, and α-tubulin. Quantitative data showing the relative protein levels of L1CAM (**C**), N-Cad (**D**), and vimentin (**E**) normalized to α-tubulin. All the experiments were repeated with at least with three biological replicates (n=3). (**F**) Representative brightfield microscope images of phagokinetic tracks created by control and KO α-syn cells on colloidal gold-coated wells were acquired using a light microscope with a 10x objective. The individual phagokinetic tracks represented by black arrow marks were measured using ImageJ software and represented in (**G**) At least 33 tracks per experimental condition were randomly chosen for quantification. A total of 100 tracks for n=3 per experimental group were used for statistical analysis. Values are mean ± s.d. *P*-values determined by a one-way ANOVA, Dunnett post hoc test.

The plot of the quantified area of single cell tracks showed an 80% decrease in motility (*P*<0.0001) for each KO clone (Fig. 4 G).

### Expressing α-syn in SH-SY5Y increases L1CAM and motility

Given that the loss of α-syn expression decreases L1CAM, N-cadherin and cell motility in two melanoma cell lines, we asked whether the reverse would be true: Does expressing α-syn in cells that lack α-syn expression increase the expression of pro-oncogenic adhesion proteins and concomitantly increase cell motility? To test this idea, we used the human neuroblastoma cell line SH-SY5Y, which has no detectable α-syn, and SH-SY5Y cells (SH/+αS) that stably express wild-type α-syn (Table 1). L1CAM, N-cadherin, vimentin, α-syn, and GAPDH levels in the cell extracts were probed by Western blotting (Fig. 5A; Fig. S5, uncropped blots), yielding the following results. First, α-syn was robustly expressed in the SH/+αS cells but not in the parental line (Fig 5A, panel 4). Second, the level of L1CAM level was 54% (*P*=0.044) higher in the SH/+αS cells than in the parental cells (Fig. 5A, panel 1; Fig. 5B). Third, N-cadherin levels were similar in the two cell lines (*P*=0.508) (Fig. 5A, panel 2; Fig. 5B). Fourth, the level of vimentin was 415% (*P*=0.001) higher in the SH/+αS cells than in the parental cells (Fig. 5A, panel 3; Fig. 5C).

**Figure 5.**
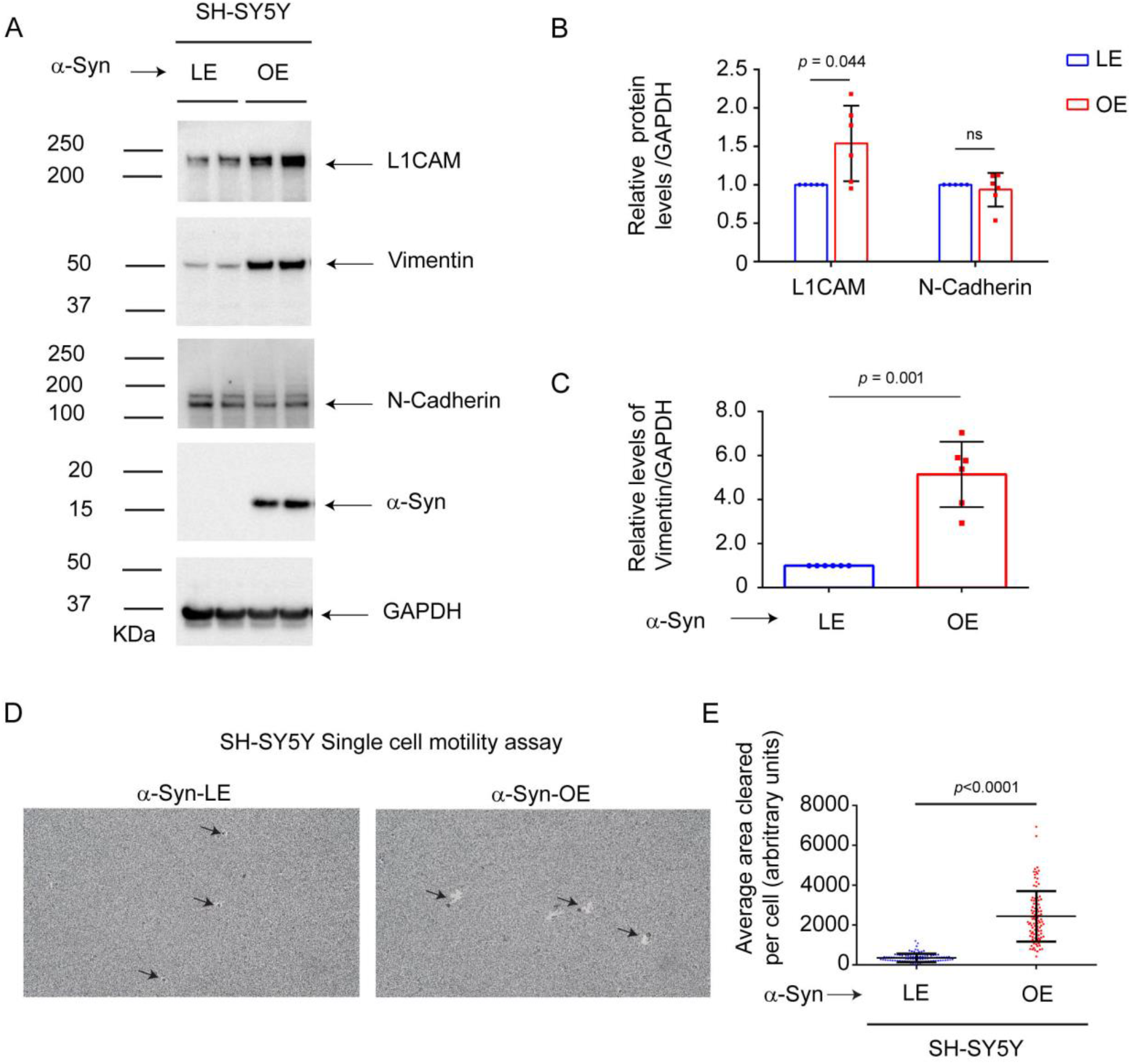
Expressing α-syn in SH-SY5Y cells increases L1CAM and motility. **(A**) Representative western blots of L1CAM, vimentin, N-Cad, α-syn, and α-tubulin in lysates of control and SH/aS cells cultured *in vitro*. Quantitative analyses representing the relative protein levels were measured by normalizing the band intensities of L1CAM and N-Cad (**B**) and vimentin (**C**) to α-tubulin. All the experiments were repeated with at least with three biological replicates (n=6). (**D**) Representative brightfield microscope images of phagokinetic tracks created by control and α-syn overexpressing SH-SY5Y cells on colloidal gold-coated wells. The images were acquired using a light microscope with a 10x objective. The individual phagokinetic tracks represented by black arrow marks were measured using ImageJ software and represented in (**E**). At least 33 tracks per experimental condition were randomly chosen for quantification. A total of 100 tracks for n=3 per experimental group were used for statistical analysis. Values are mean ± s.d. *P*-values determined by a Student’s t test.

The single-cell motility of the two SH-SY5Y cell lines was also measured. SH/+αS cells exhibited robust single-cell motility during the 24 h incubation period compared to the parental cells (Fig. 5D). The plot of the quantified area of single cell tracks shows that expressing α-syn in the SH-SY5Y cells significantly increased the single cell motility of such cells compared to control cells. Strikingly, the SH/+αS cells exhibited a 597% (*P*<0.0001) increase in single-cell motility compared to the parental cells (Fig 5E).

## Discussion

The major findings in this report are that (i) loss of α-syn expression in two human cutaneous melanoma cell lines significantly decreases L1CAM, N-cadherin, and cell motility (Figs. 1B, E, G, 4A, C, G). (ii) Loss of α-syn expression stimulates the degradation of L1CAM in the lysosome (Fig. 3). (iii) An increase in α-syn expression in SH-SY5Y cells increases L1CAM and cell motility (Fig. 5A, B, D, E)).

L1CAM is a 200-220 kDa transmembrane glycoprotein that is a member of the immunoglobulin superfamily that has a myriad of activities in the adult nervous system, including neurite outgrowth, migration, adhesion, and neuronal differentiation (for reviews see ^42-44^). L1CAM has been implicated in synaptic plasticity^42^, and, curiously, α-syn has been implicated in synaptic plasticity^38^. L1CAM, which is upregulated in several cancers of neuroectodermal and neural crest origin^45^, including melanomas, is recognized as a tumor antigen involved in motility^39-41^. Ernst et al. found that knocking down L1CAM significantly reduces metastasis in a xenograft model of human melanoma^46^. We found here that knocking out *SNCA* significantly reduces the level of L1CAM relative to control cells that express α-syn, and the decreased level of this adhesion protein likely contributes to the reduction in cell motility in two melanoma cell lines and one neuroblastoma cell line. We propose that α-syn promotes the intracellular trafficking of L1CAM (and TfR1), and that the loss of α-syn expression disrupts this trafficking, resulting in L1CAM (and TfR1) transiting to the lysosome by default for degradation (Fig. 3). The pathways affected may be sorting nexin/retromer mediated retrograde trafficking^47,48^ and/or rapid recycling^49^.

Turriani and colleagues^28^ recently suggested that “there is an inverse molecular link between PD and melanoma and that proteins that are “detrimental players” in PD are “beneficial players” in melanoma because their functions confer significant survival benefits to primary and metastatic melanoma.” Turriani showed that the compound anle128b, which disrupts the prion-like oligomers of α-syn^28^, protects neurons from α-syn-induced cell death, but in contrast, this compound promotes massive cell death of the WM983-B melanoma cell line. Treating WM983-B cells with anle138b resulted in reduced levels of aggregated α-syn, morphological changes, and disruptions in the mitochondrial membrane potential and autophagy. It was concluded that dissolving aggregated, prion-like forms of α-syn dysregulates autophagy, which suggests that these unusual, aggregated forms of α-syn promote autophagy. Certainly, there is a similarity in our work and Turriani’s in that each group proposes that synuclein is pro-oncogenic because of its role in the endolysosomal system.

E- and N-cadherin are Ca^++^-dependent cell-cell adhesion transmembrane glycoproteins^50^. The conserved cytoplasmic tails of these two proteins interact with networks of proteins involved in different cell signaling pathways. The ectodomain of E-cadherin forms homotypic dimers with neighboring cells. N-cadherin is similar in structure to E-cadherin, but its cytoplasmic tail interacts with a different set of proteins^51^. One of the hallmarks of the EMT is the upregulation of N-cadherin followed by the downregulation of E-cadherin^51^. In our hands, knocking out α-syn expression in the two melanoma cell lines significantly decreased the level of N-cadherin but not E-cadherin (Figs. 1A, C, E, 4A, C, D). It is as if loss of α-syn expression partially reverses an ‘EMT-like’ phenomenon.

Renal tubular epithelial cells, conditional knockout mice, and clinical samples of human renal tissue were recently used to assess the role of renal tubular epithelial α-syn in kidney fibrosis^52^. Bozic and colleagues found that treating HK-2 renal cells with TGF-β1, which mediates fibrosis signaling in renal epithelial cells, changed the epithelial phenotype (loss of cobblestone morphology), decreased E-cadherin, and increased α-SMA and vimentin. TGF-β1 concomitantly induced a dose-dependent decrease in *SNCA* mRNA and α-syn expression. Given the co-occurrence of the loss of the epithelial phenotype and the dysregulation of α-syn expression, the authors hypothesized that α-syn has a role in maintaining the epithelial phenotype of renal proximal tubular epithelial cells (RPTECs) in vitro. This hypothesis was supported by their discovery that overexpressing α-syn in HK-2 cells inhibited TGF-β1-induced increases in α-SMA and vimentin in vitro. They went on to show that α-syn modulates the activation of ERK1/2, Akt and p38 in vitro, that TGF-β1decreases α-syn expression via activation of the MAPK-p38 axis, and that loss of α-syn accelerates profibrotic gene expression. Our group had earlier shown that α-syn inhibits the stress-induced phosphorylation of p38 (and JNK and c-Jun) in SH-SY5Y cells^53^. Their conclusion was that α-syn plays a role in maintaining the epithelial phenotype of RPTECs. Of course, it is hard to compare our results to those of Bozic because we used non-epithelial-derived cancer cells, whereas Bozic used non-dividing renal epithelial cells. On the other hand, these two studies reveal the Dr. Jekyll and Mr. Hyde-like nature of α-syn: α-syn supports an epithelial phenotype of renal cells, whereas it supports a ‘mesenchymal-like’ phenotype of melanoma and neuroblastoma cells.

N-cadherin promotes angiogenesis^54^. In our previous study, where SK-MEL-28 control and *SNCA*-KO cells were injected into the flanks of nude mice, the tumors that formed from the *SNCA*-KO clones were significantly smaller and whiter than the tumors from control cells^30^. This lack of coloration may indicate a lack of vascularization of the tumors derived from *SNCA*-KO cells. Perhaps this lack of apparent vascularization for the tumors from *SNCA*-KO cells was due to less expression of N-cadherin.

In sum, we have shown that loss of α-syn expression in two human cutaneous melanoma cell lines results in significant decreases in two adhesion proteins, L1CAM and N-cadherin, and concomitant significant decreases in motility. We propose that α-syn is pro-survival to melanoma (and likely neuroblastoma) because it promotes the efficient trafficking of L1CAM, which in turn promotes invasion, migration, and motility.

## Materials and Methods

### Cell lines and cell culture

SK-MEL-28 and SK-MEL-29 cells were purchased from American Type Culture Collection (ATCC, Manassas, VA) and from Sloan-Kettering Memorial Center, respectively, and propagated in DMEM supplemented with 10% fetal bovine serum (FBS) and 1% penicillin–streptomycin. The human neuroblastoma cell line SH-SY5Y over expressing α-syn (SH/αS) and control SH-SY5Y cells were a kind gift of Dr. Joseph R Mazzulli (Northwestern University) and propagated in Opti-MEM supplemented with 10% fetal bovine serum (FBS) and 1% penicillin–streptomycin. CRISPR/Cas9 genome editing was used to target *SNCA* in SK-MEL-29 cells as described previously ^30^ for SK-MEL-28 cells using α-syn CRISPR/Cas9 knockout plasmid (Santa Cruz Biotechnology # sc-417273-NIC). Lentivirus particles expressing human α-syn under cytomegalovirus (CMV) promoter (Applied Biological Materials, Inc, Canada) was used to re-express α-syn in SK-MEL-28 KO cells as described previously^30^. For proteasome and autophagy inhibition experiments cells were treated with 10 μM MG132 (Thermofisher Scientific, # M7449) for 6 h and 50 nM bafilomycin A1 (Millipore, # B1793) for 5 h respectively in growth medium. The cell lines were authenticated and tested for mycoplasma contamination using MycoAlert® Mycoplasma Detection Kit (# LT07-318).

### Western blotting

Preparation of cell lysates, SDS/PAGE, and western blot analysis was carried out as described previously^30^. Briefly, the cells were lysed in RIPA lysis buffer (50 mM Tris HCl, pH 7.4, 1% NP-40, 0.5% sodium deoxycholate, 0.1% SDS, 5 mM EDTA). The lysates were centrifuged (13,000 rpm/30 min/4°C), and the protein concentrations of the supernatants were determined using DC™ Protein Assay Kit (Bio-Rad #5 000 112). Equal concentrations of protein (30 μg) from each sample were reduced with Dithiothreitol (Invitrogen™ NuPAGE™ Sample Reducing Agent # NP0004) and boiled for 10 min at 70°C in NuPAGE™ LDS Sample Buffer (#NP000). Proteins were separated by sodium dodecyl sulfate Bis-Tris polyacrylamide gel electrophoresis (SDS-PAGE) (NuPAGE™ 4 to 12%, Bis-Tris precast polyacrylamide gel, Invitrogen # NP0323BOX) and afterward transferred to polyvinylidene difluoride (PVDF) membrane using (Trans-Blot® Turbo™ Mini PVDF Transfer Pack, Bio-Rad # 1704156). After blocking the membranes with 5% blotto (G-Biosciences Blot-Quikblocker # 786-011) in phosphate-buffered saline containing 0.1% (v/v) Tween-20 (PBST) for 1 h at room temperature, the membranes were often cut into strips and the individual strips were hybridized with indicated primary antibodies overnight at 4°C followed by incubation with respective horse radish peroxidase (HRP) conjugates.

The immunoreactive bands were visualized using an enhanced chemiluminescence substrate (Clarity™ Western ECL Substrate, Bio-Rad #170-5060) and the images were acquired using Biorad Chemidoc-MP imaging system. The band intensities of proteins of interest were quantified and represented as relative protein levels normalized to housekeeping protein α-tubulin using ImageJ software. Antibodies (with dilutions) are given in Table 2.

**Table 2.**
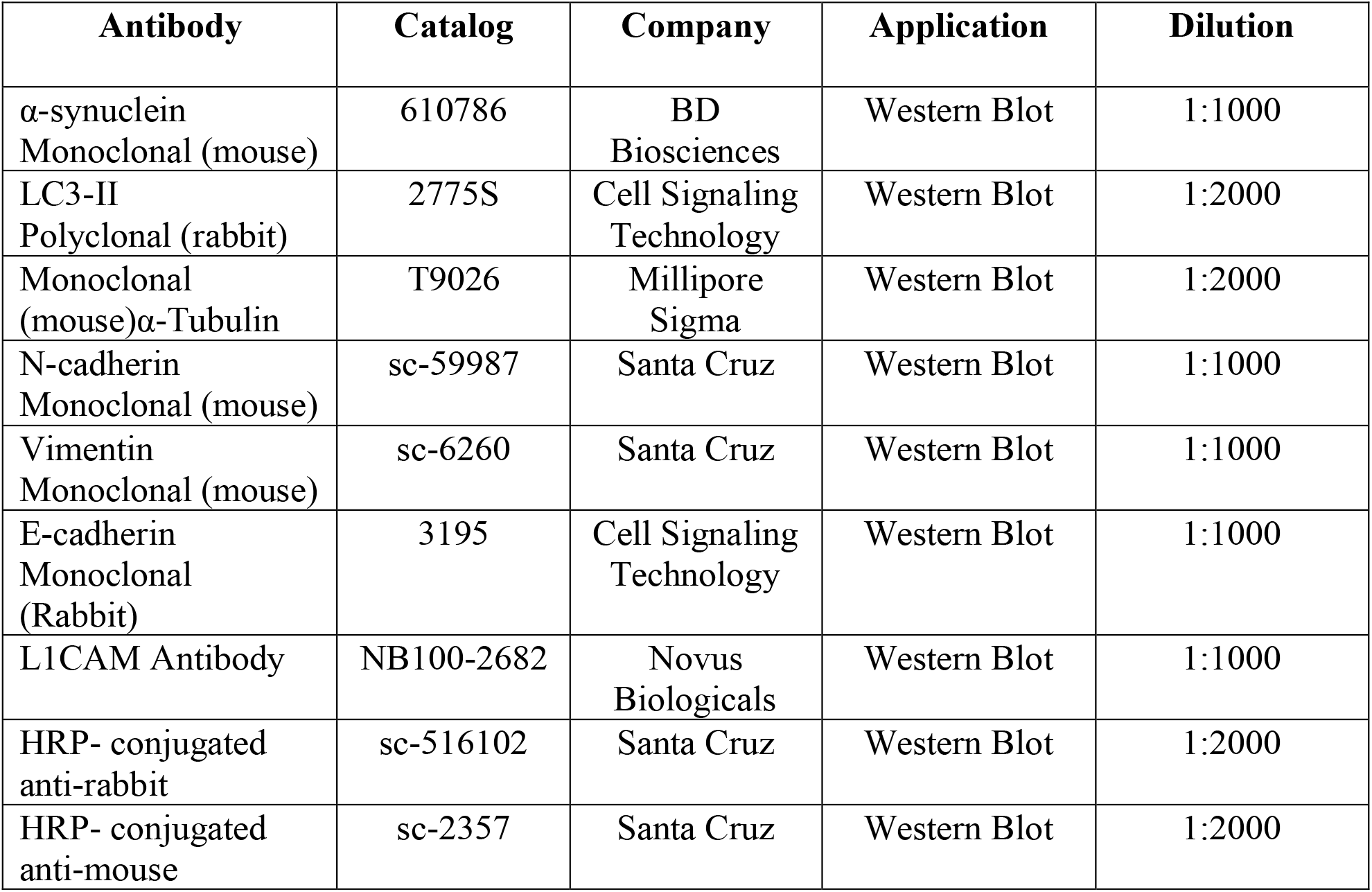

### Migration and invasion assays

The invasion and metastatic potential of the cell lines in Table 1 was determined by transwell assays using Boyden chambers with an 8-μm pore size (Costar; Corning, Inc. # CL3464). Briefly, the cells were suspended in serum-free DMEM and were seeded onto the apical chamber with (invasion assay) and without (migration assay) matrigel (Corning, Inc. #356234), and 750 μl complete medium containing 10% FBS was added to the lower chamber of the transwell for 24 h. After incubation at 37 °C for 24 h, the cells migrated/ invaded through the lower surface and were fixed with 4% paraformaldehyde for 5 min and then stained with 0.1% crystal violet for 10 min to allow the cells to be visualized. The cells in the upper chamber of the transwell were carefully removed with a cotton swab, and the images of cells in the lower chamber of the transwell were taken using Olympus inverted light microscope.

### Phagokinetic single-cell motility assay

The motility of control and *SNCA*-KO melanoma cell lines was measured by tracking the ability of cells to clear gold from their path. For this, 6-well plates were coated with 2 ml of 1% BSA, incubated for 3 h in a humidified CO_2_ incubator at 37 °C, and washed with absolute ethanol. The wells were then coated with a homogenous layer of colloidal gold solution, prepared as follows: A total of 3.85 mL of sterile H_2_O, 630 μL of 14.5 mM AuHCl4, and 2.1 ml of 36.5 mM Na_2_CO_3_, followed by boiling at 100 °C for 5 min and addition of 0.1% of formaldehyde. The colloidal gold-coated plates were incubated in CO_2_ incubator at 37 °C for 24 h, and cells were seeded at a final number 1 × 10^3^ cells per well in a complete medium containing 10% FBS. After 24 hours, the wells were imaged using Olympus inverted light microscope, and the tracks were quantified using ImageJ Software.

### RNA extraction, cDNA preparation, and qPCR

Total RNA was extracted from cells using E.Z.N.A column-based total RNA kit (Omega BioTek) following the manufacturer’s instructions. The concentration and quality of the extracted RNA were determined on a NanoDrop spectrophotometer (Thermo Scientific). cDNA was synthesized from total purified RNA (1 μg) from each sample by using iScript cDNA synthesis kit (Bio-Rad) according to the manufacturer’s protocol. qPCR was performed using Applied Biosystems TaqMan™ Gene Expression Assays with primer/probe sets for *SNCA* (Hs00240906), *L1CAM* (Hs01109748), *GAPDH* (Hs02786624). ΔΔCT method was adopted to calculate the relative amount of each mRNA normalized to the housekeeping gene GAPDH. Data were analyzed using the comparative CT method, and the fold change was calculated using the 2^−ΔΔCT^ method using Bio-Rad CFX384 Touch Real-Time PCR System and software (Bio-Rad). The results were expressed either as the relative log_2_ FC (fold change) relative values.

### Statistical analyses

Hypothesis testing method included a one-way analysis of variance (ANOVA) with a Dunnett post hoc test when comparing multiple groups to control or two tailed Student’s t-test when comparing two groups. All data were analyzed using GraphPad Prism (version 6) software. All values were expressed as mean ± standard deviation (s.d.) of at least three independent experiments (biological replicates). *P*-value of < 0.05 was considered significant.

## Acknowledgments

This study was supported by funds from the Feist-Weiller Cancer Center and the Chancellor of LSU Health Sciences Center in Shreveport to S.N.W.

## Author Contributions

N.G and S.R. contributed equally to the study’s conception, experimentation, data analysis, interpretation, and revision. S.N.W. contributed to the study’s conception, supervised the study, data analysis, and wrote and edited the manuscript. All authors read and approved the final manuscript.

## Data availability

All data from this study are contained within the article and its Supplementary Information.

