## Supplemental Figures for "Knocking out alpha-synuclein in melanoma cells downregulates L1CAM and decreases motility"

#, equal contributions

**Figure S1**

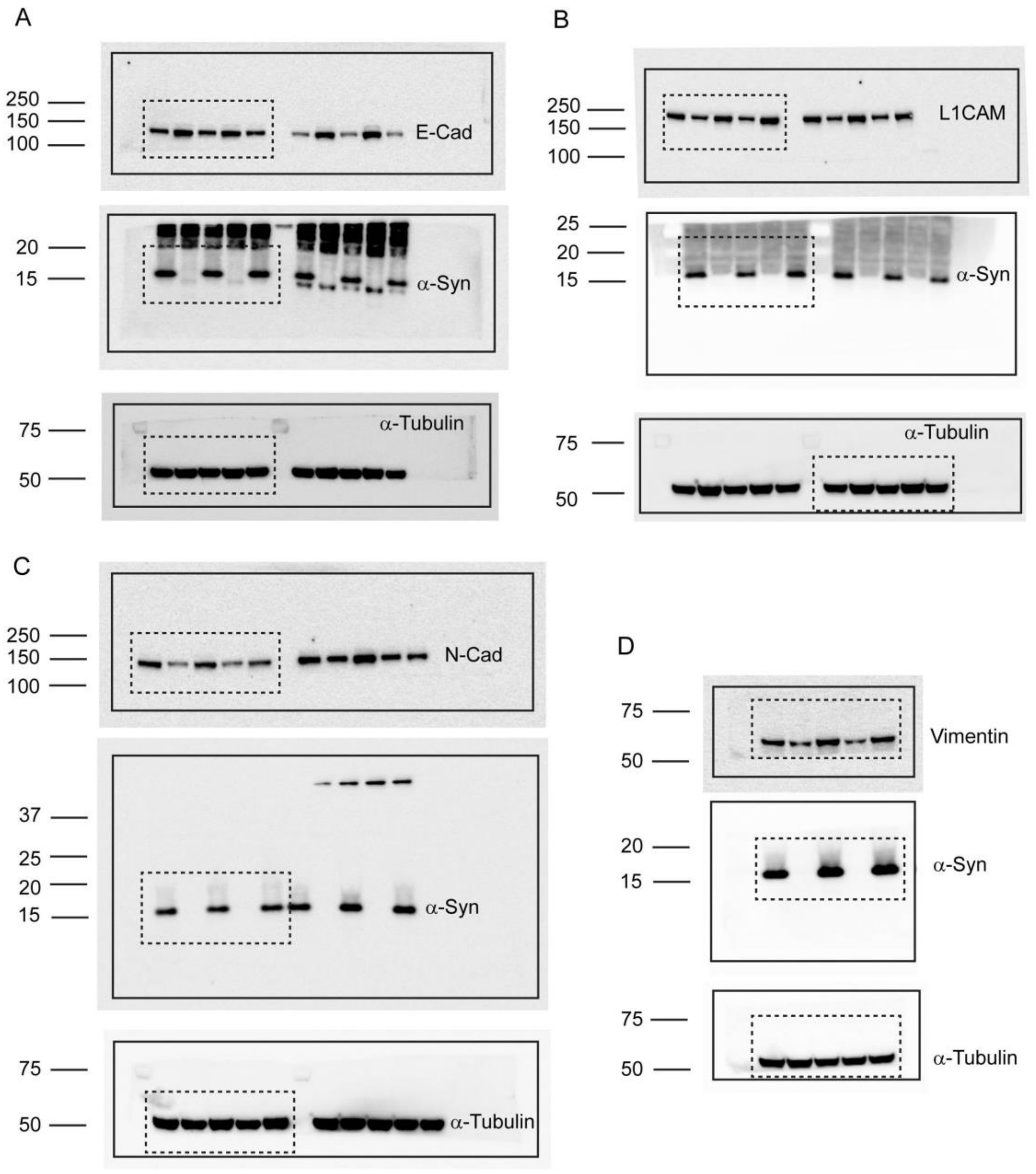


**Supplementary Figure S1**

(**A-D**) Full-length original uncontrasted Western blots are shown cropped in Figure 1 A, B, C, and D. The edges of the blots were represented by a solid rectangle, and the cropped area was represented by a broken rectangle.

**Figure S2**

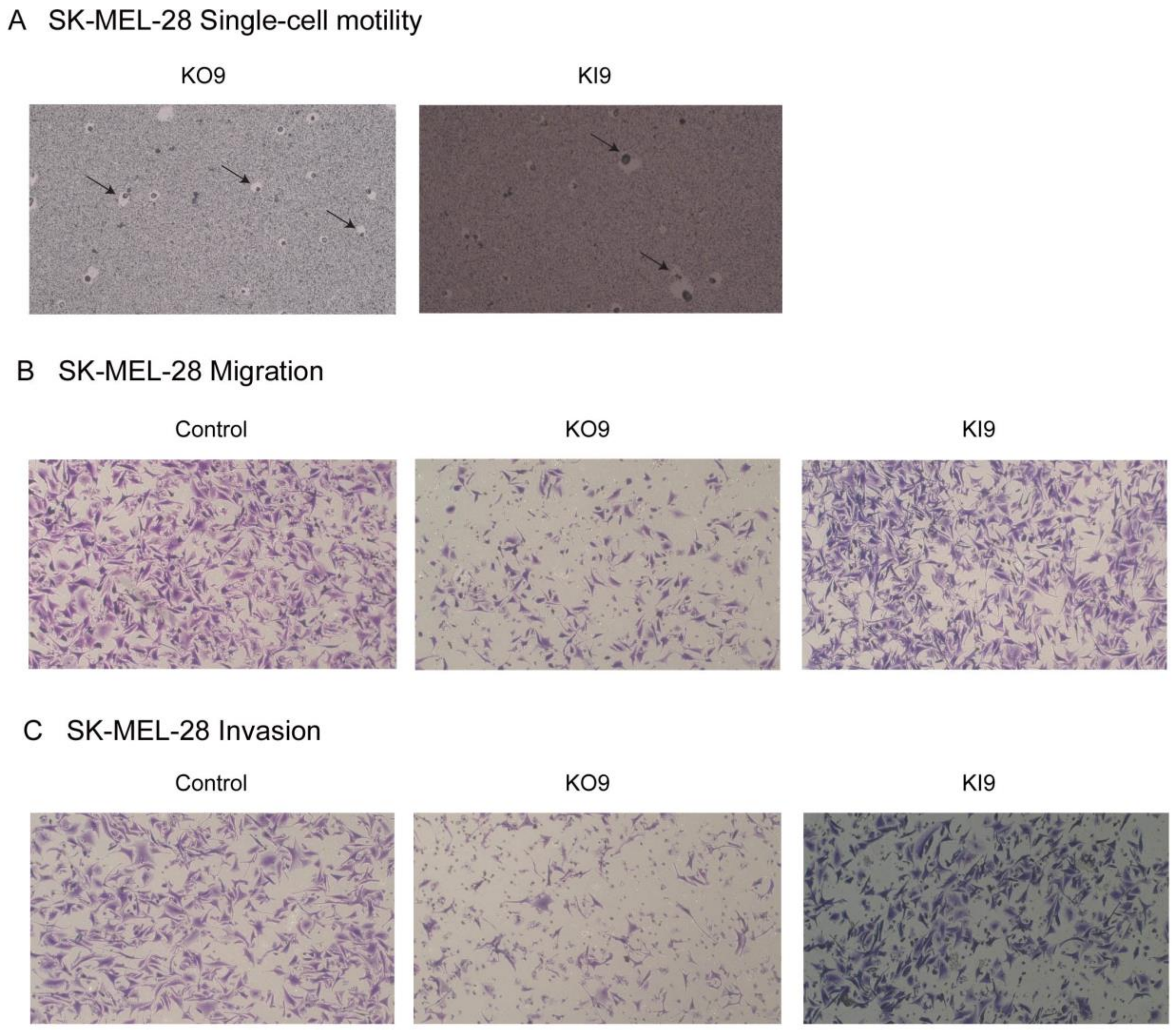


**Supplementary Figure S2**

(**A**) Representative light microscope images of phagokinetic tracks created by KO9 and KI9 cells on colloidal gold-coated wells for the data shown in Figure 1E and F. Black arrows mark individual phagokinetic tracks in respective cell lines. (**B**) Representative images (10X objective) of migrated control, KO9, and KI9 cells (after 24 hours) for the data shown in Figure 2A. (**C**) Representative images (10X objective) of invaded control, KO9 and KI9 cells (after 24 hours) per 10X field for the data shown in Figure 2B.

**Figure S3**

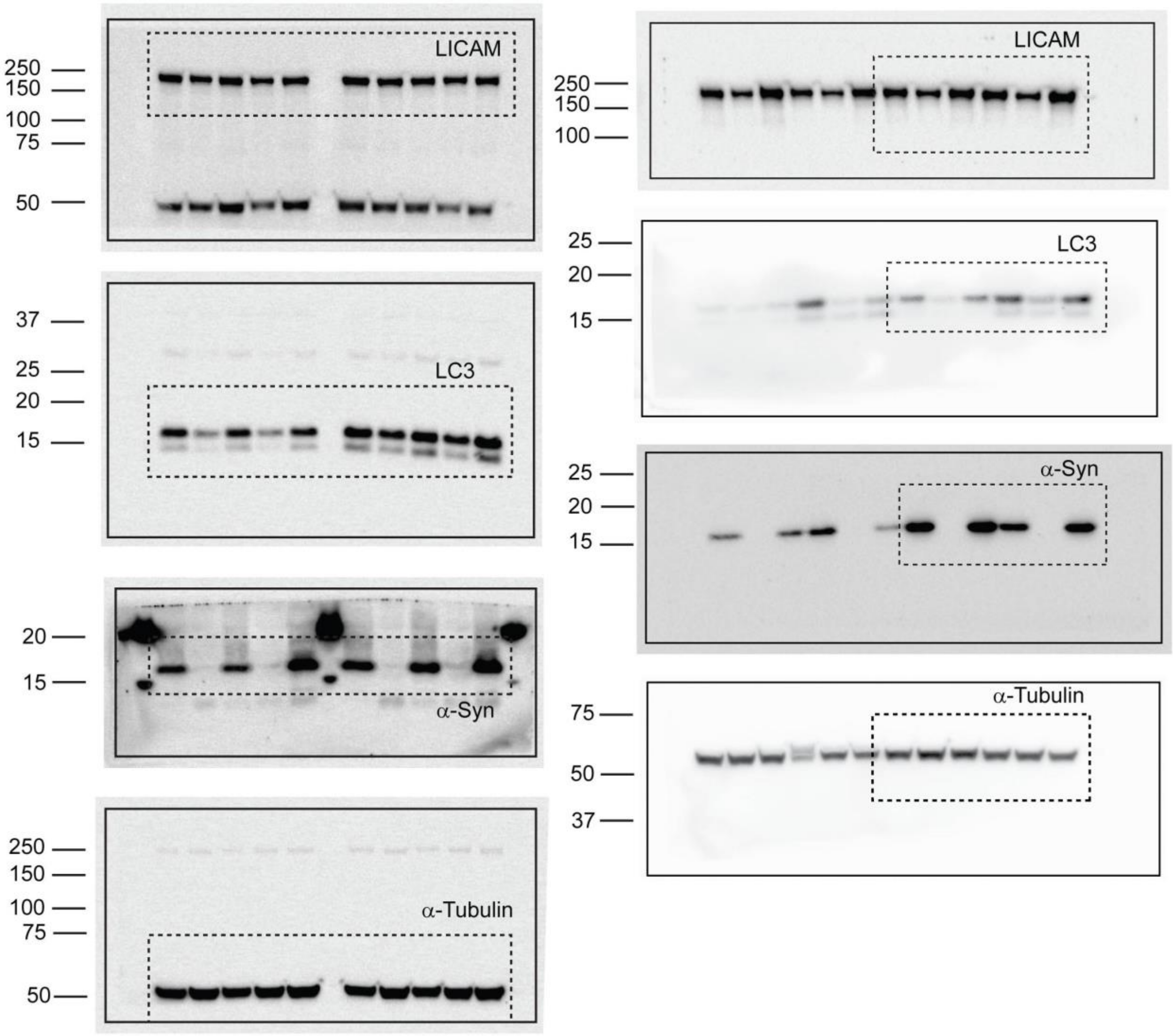


**Supplementary Figure S3.**

Full-length, original uncontrasted Western blots are shown cropped in Figures 3 A and C. A solid rectangle represented the edges of the blots and the cropped area was represented by a broken rectangle.

**Figure S4**

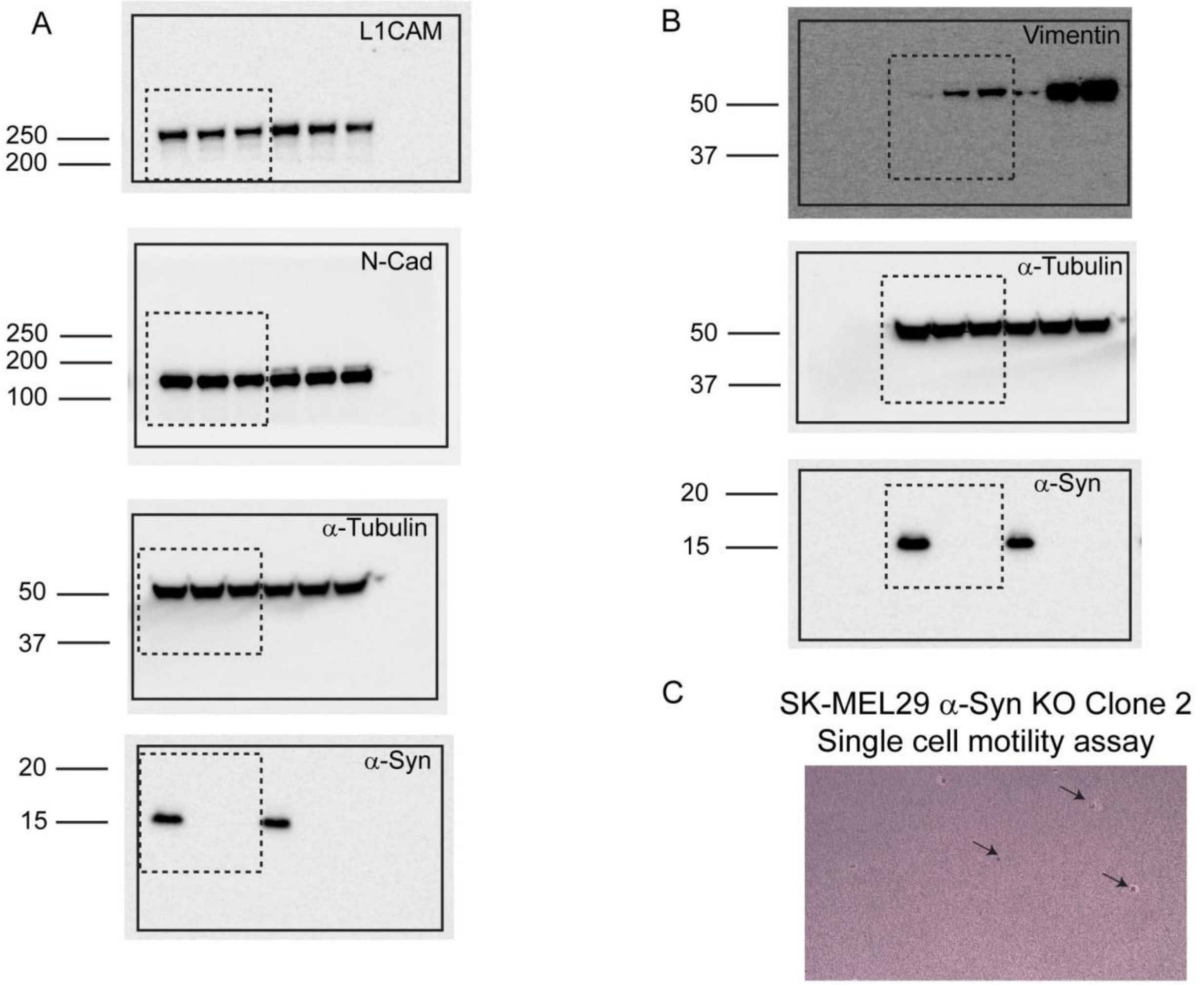


**Supplementary Figure S4**

(**A**, **B**) Full-length, original uncontrasted Western blots which are shown cropped in Figures 4 A and B. The edges of the blots were represented by a solid rectangle and the cropped area was represented by a broken rectangle. (**C**) Representative light microscope images of phagokinetic tracks created by SK-MEL29 α-syn KO Clone 2 on colloidal gold-coated wells for the data shown in Figure 4F and G. Black arrows mark individual phagokinetic tracks in respective cell lines.

**Figure S5**

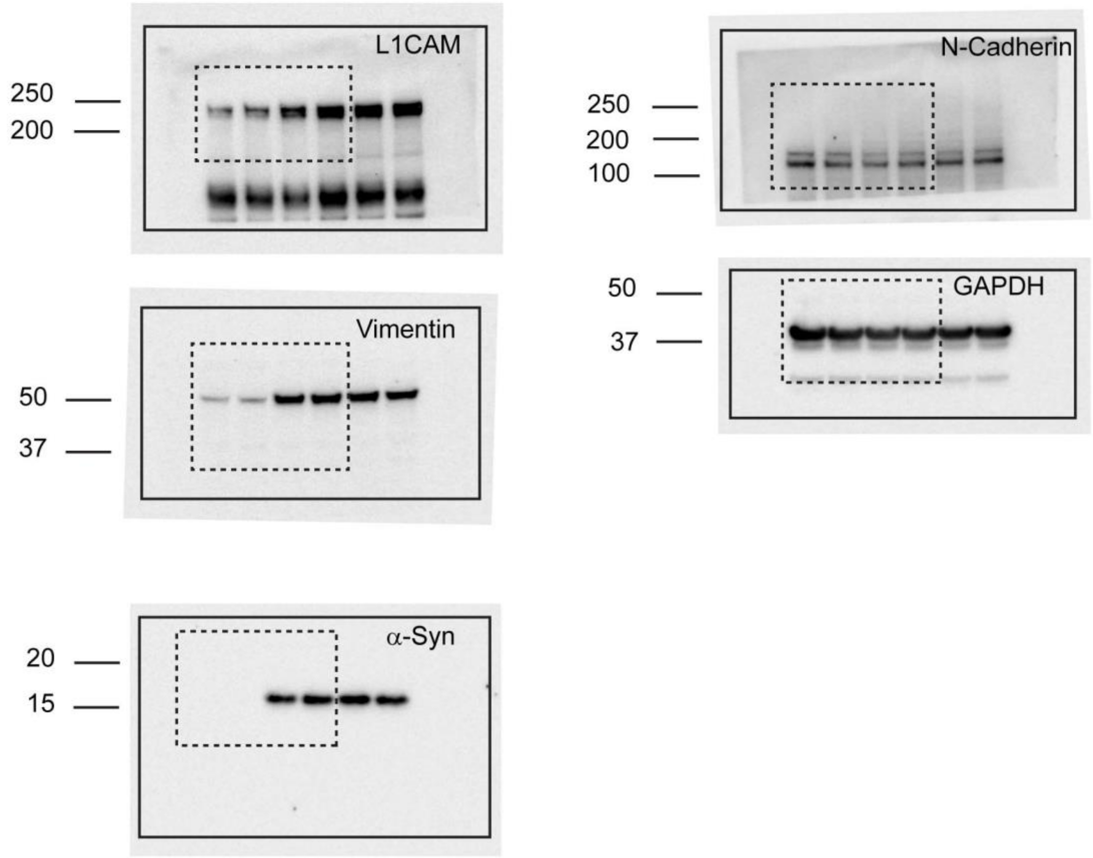


**Supplementary Figure S5**

Full-length, original uncontrasted Western blots are shown cropped in Figure 5A. The edges of the blots were represented by a solid rectangle and the cropped area was represented by a broken rectangle.
